# Disruptions in White Matter Maturation and Mediation of Cognitive Development in Youth on the Psychosis Spectrum

**DOI:** 10.1101/423574

**Authors:** Catherine E. Hegarty, Dietsje D. Jolles, Eva Mennigen, Maria Jalbrzikowski, Carrie E. Bearden, Katherine H. Karlsgodt

**Author notes:** **Corresponding Author Information**: Katherine H. Karlsgodt, Ph.D., Assistant Professor in Psychology and Psychiatry, 1285 Franz Hall Box 951563, University of California, Los Angeles, Los Angeles, CA 90095. Disclosures*: The authors have no conflicts of interest or financial interests to disclose.

## Abstract

**Background:** Psychosis onset typically occurs in adolescence, and subclinical psychotic experiences peak in adolescence as well. Adolescence is also a time of critical neural and cognitive maturation. Using cross-sectional data from the Philadelphia Neurodevelopmental Cohort, we examine whether regional white matter (WM) development is disrupted in psychosis spectrum (PS) youth whether WM maturation mediates the relationship between age and cognition in typically developing (TD) and PS youth. A third group with intermediate symptom severity (limited PS [LPS]) was included in follow-up analyses to determine whether age-related disruptions in WM scaled with symptom severity.

**Methods:** We examined WM microstructure, as assessed via diffusion tensor imaging, in 707 individuals (aged 10–22 years; 499 TD, 171 PS, 37 LPS) by using Tract-Based Spatial Statistics. Multiple regressions were used to evaluate age x group interactions on regional WM indices. Mediation analyses were conducted using a bootstrapping approach.

**Results:** There were age x group interactions on fractional anisotropy (FA) in the superior longitudinal fasciculus (SLF) and retrolenticular internal capsule (RLIC). SLF FA mediated the relationship between age and Complex Cognition in TD, but not PS. Further, inclusion of LPS youth showed that the relationship between age and SLF FA decreased with increasing symptom severity

**Conclusions:** Our results show aberrant age-related changes in SLF and RLIC FA in PS youth. SLF development supports emergence of specific higher-order cognitive functions in TD youth, but not in PS. Future mechanistic explanations for these relationships could facilitate development of earlier and refined targets for therapeutic interventions.

## Introduction

Schizophrenia (SZ) has a lifetime prevalence of 0.75% [1], but — consistent with our understanding of a psychosis continuum [2, 3]— a much greater percentage of the population (5-8%) endorse the presence of subclinical psychotic experiences [4, 5]. These experiences are qualitatively similar to the hallmark psychotic symptoms of SZ, but have decreased severity and frequency. Although below the clinical threshold, these symptoms are still distressing and are associated with lower reported happiness [6], increased suicidality [7, 8], and significant occupational and social impairments [9]. Subclinical psychotic symptoms also share etiological risk factors, cognitive correlates, and symptom profiles with SZ [3], and can be a harbinger of subsequent full-blown illness. Similar to transition rates for clinical high-risk youth [10], up to 25% of individuals who report these experiences convert to a diagnosed psychotic disorder by adulthood [11, 12]. For those who do not convert, approximately 30-40% continue to experience these subclinical symptoms into early adulthood [13, 14] and, potentially, throughout life [15].

The onset period for SZ begins in late adolescence and, similarly, subclinical psychotic symptoms peak during adolescence [3], coinciding with critical periods of dynamic change in brain connectivity and cognition. One possible explanation for this age-related component of psychosis is a disrupted neurodevelopmental trajectory of white matter (WM) maturation. During typical development, WM volume linearly increases from childhood until early adulthood [16, 17], a pattern which is thought to reflect axon growth, organization of axons into bundles, and myelination [18]. WM integrity is putatively reflected by the principal diffusion tensor imaging (DTI) measure, fractional anisotropy (FA)[19]. Studies evaluating WM indices have shown regionally specific age-related changes with the overarching trend of FA increasing into adulthood [20, 21]. Although both early-onset and chronic SZ have been associated with reduced FA, particularly in frontal and temporal regions (for review see [22, 23]), the manner in which the developmental trajectories of regional structural connectivity in SZ diverge from those in typically developing (TD) youth is unknown. Here, we propose to use TD and psychosis spectrum (PS) groups from the Philadelphia Neurodevelopmental Cohort (PNC), to explore how psychosis affects age-associated regional changes in WM. While one previous study reported an age by group interaction on regional grey matter volume in this cohort [24], no studies have yet investigated such interactions of WM indices as assessed by DTI.

Subclinical PS symptoms, as with SZ symptoms, have been associated with cognitive deficits, both generally and in specific domains [25-29]. However, despite the demonstrated link between psychosis and cognition, and the known relationship of WM to cognitive development [30-32], it is not yet established whether regional WM measures mediate the relationship between age and cognitive deficits associated with psychosis. To address these questions, we first evaluated whether the age-related pattern of regional WM development was disrupted in youth with elevated subclinical psychotic symptoms. Then, to account for the dimensional nature of PS, we extended significant interaction findings with follow-up analyses including a third group, limited psychosis (LPS), whose endorsement of subclinical symptoms is intermediate to the TD and PS groups. Finally, we evaluated whether any implicated tracts mediated the relationship between age and performance in four cognitive domains, and whether that relationship differed between TD and PS youth.

## Methods

### Participants

The PNC is a publicly available population-based sample of more than 9500 individuals from the greater Philadelphia area (ages 8-22; for description see [33]). All subjects provided medical history, clinical and cognitive data. A subset of 1445 participants underwent neuroimaging, with DTI scans acquired for 1312 individuals. All data presented in this manuscript were collected at The University of Pennsylvania (Penn) and analyzed at University of California, Los Angeles. Psychopathology self- and parent-report ratings and medical histories were obtained from the GOASSESS structured computerized instrument.

In addition to previously described PNC exclusion criteria for medical problems impacting brain function [33], participants were excluded if they had more than 2.5mm of total Euclidian distance movement during the DTI scan (n=44), had a history or current diagnosis of autism spectrum disorders (n=13) or were indicated as having other neurological or non-psychotic psychiatric diagnoses (n=452; See Supplementary Information). Due to the limited number of younger PS individuals, participants under the age of 10 years (n=122) were also excluded to ensure groups were matched for age distribution. After excluding participants who met one or more of the aforementioned criteria, 707 individuals remained for analysis.

### Subclinical Grouping

Psychosis screening and classification of individuals into TD, PS and LPS groups were performed as described by Calkins [34] and are detailed in Supplementary Information. Our primary analyses consisted of PS and TD groups. Given the small size of the LPS sample (n=37), these individuals were only included as a third group in follow-up interaction analyses to allow us to do a preliminary investigation of more subtle dimensional changes.

### Cognitive Data

Cognitive testing was administered via the Penn Computerized Neurocognitive Battery (CNB). Descriptions of the metrics and calculation of efficiency scores for Complex Cognition (language reasoning, nonverbal reasoning and spatial ability), Executive Control (mental flexibility, attention, and working memory), Episodic Memory (verbal memory, face memory, and spatial memory) and Social Cognition (emotion identification, emotion differentiation, and age differentiation) were calculated as previously described [35]. Our analyses focused on efficiency scores — obtained via the PNC data release and based on the entire neurocognitive sample (n=9138) — which reflect the sum of the z-scores for accuracy and speed. Analyses of covariance (ANCOVAs), with age and sex as covariates, were used to evaluate significant differences between efficiency scores of TD and PS. A Bonferroni correction for each of the four domains was applied to the significance threshold value, resulting in a threshold of p<(0.05/4)=0.0125.

### DTI Processing

The PNC protocol divides a 64-direction DWI set into two independent 32-direction sequences (b-value: 1000 S/mm^2^; see [33] for additional parameter details). The two sequences were concatenated before applying standard preprocessing using tools from FMRIB Software Library’s (FSL’s) Diffusion Toolbox. Specifically, Eddy Correct used affine registration to correct for distortions due to eddy currents and head motion. Registered b0 and concatenated files were skull stripped and masked using Brain Extraction Tool (BET). FA images were calculated by fitting a diffusion tensor model at each voxel with DTIFit. We then implemented tract based spatial statistics (TBSS; [36]) in accordance with the ENIGMA-DTI pipeline (http://enigma.ini.usc.edu/protocols/dti-protocols, [37]), and extracted regions of interest (ROIs) from the JHU White-Matter Tractography Atlas [38-40]. Bilateral ROIs were generated by averaging across both hemispheres. We tested average FA 25 total ROIs (Figure S1, Table S1).

FA, often described as an index of general WM integrity [41-43], was our principle measure of interest. Additional diffusivity measures were examined in follow-up analyses to further probe the microstructure underlying regional FA. Specifically, axial diffusivity (AD) reflects diffusivity parallel to the axon, and is thought to reflect fiber organization. Radial diffusivity (RD) is perpendicular to axons, and an indirect measure of myelination. Mean diffusivity (MD) is an average of diffusivity across all three axes [41]. AD, MD, and FA were calculated by DTIFit, while RD was computed as the average of the second and third eigenvalue images. ENIGMA-TBSS processing was also applied to these non-FA DTI measures.

### Statistical Analysis

Statistical analysis and graphical representations of data were performed in STATA (v.15, StataCorp 2017). If an individual’s FA for a given tract was more than 1.5 interquartile ranges (IQR) below the first quartile of data (Q_1_ – [1.5·IQR]) or above the third quartile (Q_3_ + [1.5·IQR]), it was deemed an outlier and excluded from subsequent analysis of that tract. The number of individuals used in analysis of each tract is reported in Table S1. To test for an age x group interaction and main effect of group on regional FA, a regression model including age, sex, and group (TD and PS) was used. The regression was followed by evaluation of marginal effects, computed at 1-year age intervals, for interaction analyses. A corrected significance threshold for all regression analyses was set at p<(0.05/25)=0.002, which corrects for the 25 tracts analyzed.

In the two tracts with significant age x group interactions on FA, we conducted a series of follow-up analyses. To explore whether the extent of subclinical symptoms affected age-associated FA changes, analyses including LPS youth as a third group were conducted to determine if an age x group (TD, LPS and PS) interaction was still present. Post-hoc contrasts then compared the slope of marginal effects between the groups to determine whether LPS represented an intermediate phenotype. We also tested for age x group interactions on hemispheric FA and non-FA measures. A significance threshold of p<(0.05/25)=0.025 was applied to account for two tracts undergoing follow-up analyses.

### Post hoc Mediation Analyses

For tracts in which a significant age x group interaction was found for FA, we conducted post-hoc mediation analyses to evaluate whether the relationship between age and any of the four cognitive domains was mediated by FA and, if so, whether that mediation differed between TD and PS youth. Individuals missing a score for a given cognitive domain were only excluded from mediation analyses for that particular domain.

To test for mediation, we used methods described by MacKinnon and colleagues [44]. Path coefficients between the independent (*X*), dependent (*Y*) and mediator (*M*) variables were determined using three regression equations (Figure1). We first determined whether initial criteria for mediation were met. Specifically, whether path coefficients *a* (effect of age on FA), *b* (effect of FA on cognition) and *c* (effect of age on cognition) were all significant (Figure 1). If these criteria were satisfied, we continued to test the significance of the indirect (i.e. mediated) effect, computed as the product of regression coefficients *a* and *b* [45].

**Figure 1.**
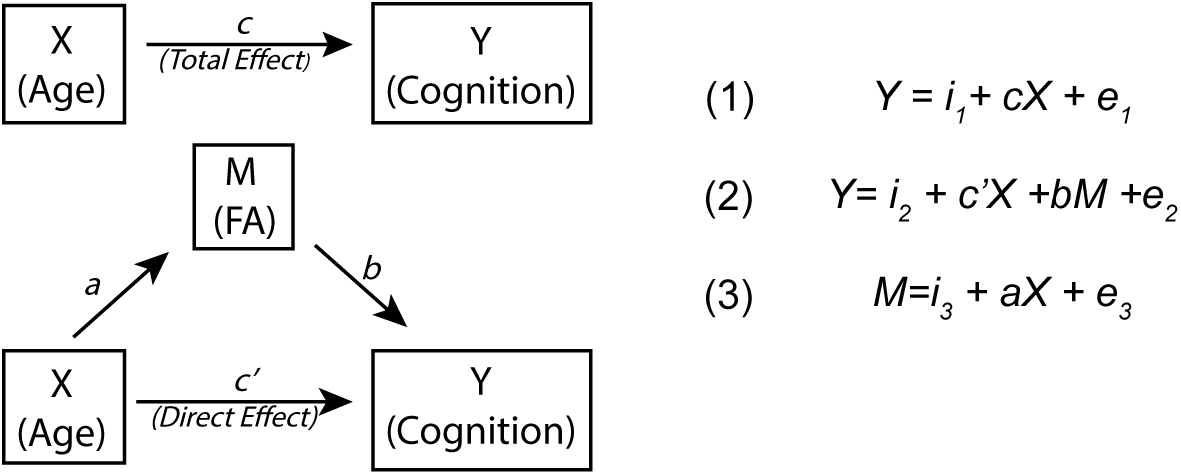
Schematic of a simple mediation model. We examined whether regional FA significantly mediated the effect of age on cognition. In a mediation model, the relationship between an independent variable (*X*) and a dependent variable (*Y*) is influenced by the (non-observable) mediator variable (*M*). Intercepts and residuals for each equation are denoted by *i* and *e*, respectively. The total effect (*c*) is the sum of both the direct (*c’*) and mediated (*ab*) effects. The total effect, c, was determined with Equation (1). Coefficients a and b were determined with Equations (2) and (3), respectively. The direct effect (*c’*) was determined with Equation (2). Mediation is determined by assessing the significance of the mediated effect (*ab*) with a bootstrapping approach. Abbreviations: FA=fractional anisotropy.

We used a bootstrapping approach, in which data were resampled with replacement 1,000 times, to assess for significance of the mediated effect. The bootstrapping approach allows for confidence intervals to be built from the resampled data without assumptions of normality. To reduce the chance of Type I errors, we increased our confidence interval to 99%. Confidence intervals that included 0 were deemed as nonsignificant. Finally, we determined the proportion of the total effect that was mediated by dividing the indirect effect (*ab*) by the total effect (*c*=*c’*+ *ab*).

## Results

### Participants

Out of 707 individuals included in the analysis, 499 were classified as TD, 171 as PS and 37 as LPS. There were no significant differences between groups in age or sex distribution (Table 1).

**Table 1.** Demographic information for participants. Abbreviations: TD=typically developing, PS = psychosis spectrum, LPS= limited psychosis spectrum, SD=standard deviation, F=female, GAF=Global Assessment of Functioning Score, yrs.=years.

|  | Group |  |  | Statistic | Significance |
| --- | --- | --- | --- | --- | --- |
|  | TD | PS | LPS |  |  |
| Age (yrs. ± SD) | 15.75 ± 3.36 | 15.83 ± 2.89 | 15.32 ± 3.13 | F(2,706)=0.371 | p=0.690 |
| TD vs PS |  |  |  | t(1,668)=-0.28 | p=0.784 |
| TD vs LPS |  |  |  | t(1,534)=0.75 | p=0.454 |
| PS vs LPS |  |  |  | t(1,206)=0.95 | p=0.342 |
| n (%F) | 499 (49.1%) | 171 (53.2%) | 37 (37.8%) | $\chi^2(2, N=707)=2.99$ | p=0.224 |
| TD vs PS | | | | $\chi^2(1, N=670)=0.864$ | p=0.353 |
| TD vs LPS | | | | $\chi^2(1, N=536)=1.75$ | p=0.186 |
| PS vs LPS | | | | $\chi^2(1, N=208)=2.88$ | p=0.09 |
| GAF | 79.81 ± 21.07 | 69.22 ± 19.7 | 73.22 ± 15.86 | F(2,654)=16.14 | p=1.44 $\times 10^{-7}$ |
| TD vs PS | | | | t(1,623)=5.53 | p=4.8 $\times 10^{-8}$ |
| TD vs LPS |  |  |  | t(1,499)=1.74 | p=0.083 |
| PS vs LPS |  |  |  | t(1,186)=-1.079 | p=0.282 |
| Education (yrs.) | 8.61 ± 3.27 | 8.13 ± 2.64 | 8.08 ± 3.06 | F(2,704)=1.797 | p=0.167 |
| TD vs PS |  |  |  | t(1,668)=1.73 | p=0.084 |
| TD vs LPS |  |  |  | t(1,534)=0.949 | p=0.343 |
| PS vs LPS |  |  |  | t(1,206)=0.952 | p=0.342 |

### DTI Analysis

Multiple regression analyses, including age, sex and group, for each of the 25 tracts revealed that only the superior longitudinal fasciculus (SLF) and retrolenticular portion of the internal capsule (RLIC) showed a significant interaction between age and group (TD vs PS; p<0.002). The SLF (Figure 2A) was the only tract in which a significant main effect of group was observed (Table S1). Additional tracts showed age x group interactions meeting the thresholds for trend-level significance (p<0.004) and uncorrected significance (p<0.05; Table S2).

**Figure 2.**
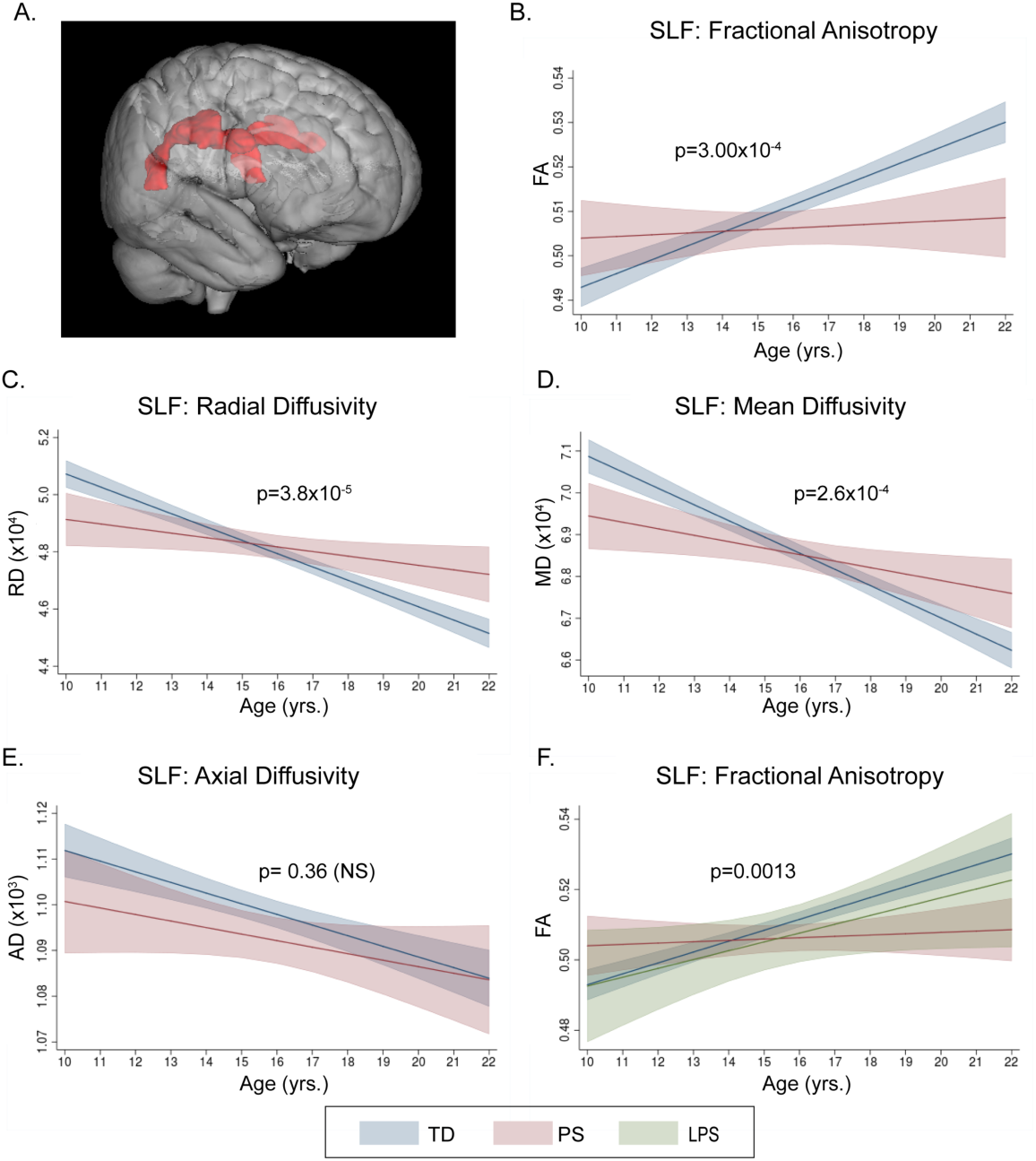
Age-associated changes in the Superior Longitudinal Fasciculus. (A) 3-dimensional representation of the SLF tract. In analyses of TD and PS groups, there was a significant(p<0.002) interaction between age and group on SLF (B) FA. Follow-up analyses revealed additional significant (p<0.025) age x group interactions on (C) RD and (D) MD, but not on (E) AD. (F) Follow-up analyses including a third group, LPS, also revealed a significant age x group interaction on SLF FA. Graphs show predicted margins with 95% confidence intervals. Significance values represent corrected p-values for age x group interactions. Abbreviations: SLF=superior longitudinal fasciculus, FA=fractional anisotropy, RD=radial diffusivity, MD= mean diffusivity, AD=axial diffusivity, TD=typically developing, PS= psychosis spectrum, LPS=limited psychosis spectrum.

#### Superior Longitudinal Fasciculus

All three variables (age, sex, group) in the regression model significantly explained SLF FA (R^2^=0.14, F(4,660)=26.86, p=1.15×10^-20^), with a significant interaction between age and group (β=-0.002, F(1,660)=13.20, p=3.0×10^-4^; Figure 2B), and each variable independently significantly contributing to the model. Evaluation of regression coefficients for each group revealed that TD showed a gradual increase in FA with age (β=0.003, p=3.68×10^-19^), while there was no significant relationship between age and FA in PS youth (β=4.9×10^-4^, p=0.48). Confirmatory analyses indicated no significant group x sex (F(1,660)=2.04, p=0.15) or age x group x sex (F(1,657)=1.18, p=0.28) interactions on SLF FA.

Secondary analyses revealed significant age x group interactions in both the right (β=-0.0032, F(1,660)=15.51, p=1.0×10^-4^) and left (β=-0.0022, F(1,660)=8.51, p=0.0036) SLF. Of the three non-FA SLF diffusivity measures, both RD (β=3.30×10^-6^, F(1,660)=17.21, p=3.8×10^-5^; Figure 2C) and MD (β=2.54×10^-6^, F(1,660)=13.46, p=2.6×10^-4^; Figure 2D) showed a significant age x group interaction. TD youth showed significant decreases in RD (TD: β=-4.47×10^-6^, p=3.8×10^-32^) and MD with age (β=-3.82×10^-6^, p=6.8×10^-32^). PS youth also showed lower RD (β=-1.36×10^-6^) and MD (β=-0.13×10^-6^) with increasing age, but this association was not significant (p>0.025). There was no significant interaction between age and group for AD (Figure 2E).

Preliminary analyses including LPS youth as a third group in the regression model also resulted in a significant age x group interaction (F(2, 695)=6.68, p=0.0013; Figure 2F). Post-hoc contrasts revealed that the slope of the relationship between age and FA for LPS youth did not significantly differ from TD (t(1,695)=0.43, p=0.664) or PS (t(1,695)=-0.144, p=0.149). Graphical representation of the interaction and evaluation of the regression coefficient between age and SLF FA in LPS revealed a positive, non-significant association between age and FA (β=0.0025).

#### Retrolenticular Portion of Internal Capsule

In the RLIC (Figure 3A), all three variables in the multiple regression analysis collectively explained FA (R^2^=0.037, F(4,657)=6.24, p=0.0001), with a significant interaction between age and group on RLIC FA (β=-0.002, F(1,657)=10.44, p=0.0013; Figure 3B), but no significant main effect of group (p>0.002). Evaluation of regression coefficients for each group indicated that TD and PS exhibited opposite patterns, with a gradual linear increase in RLIC FA across development for TD (β=0.001, p=0.001) and a non-significant decrease in PS (β=-0.001, p=0.062) with increasing age. Confirmatory analyses indicated no significant group x sex (F(1,657)=0.21, p=0.644) or age x group x sex (F(1,654)=0.17, p=0.681) interactions.

**Figure 3.**
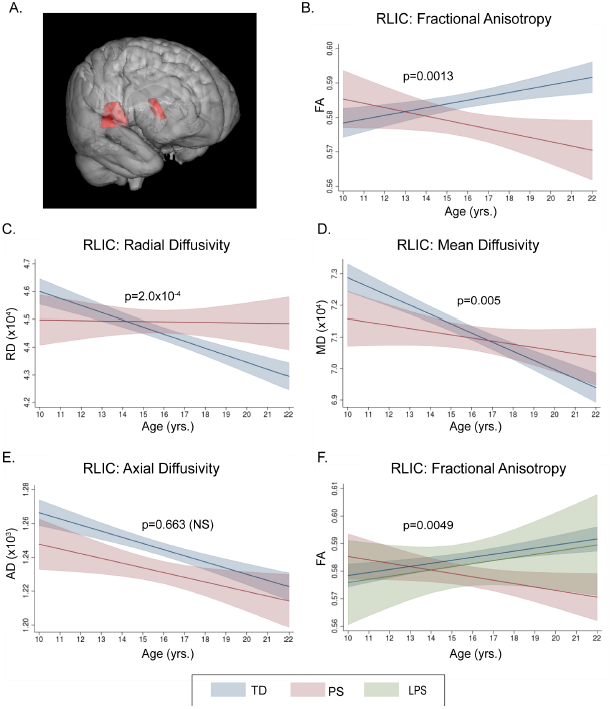
Age-associated changes in the Retrolenticular Portion of Internal Capsule. (A) 3-dimensional representation of the RLIC tract. In analyses of TD and PS groups, there was a significant interaction (p<0.002) between age and group on RLIC (B) FA. Follow-up analyses also revealed significant age x group interactions (p<0.025) on (C) RD and (D) MD, but not on (E) AD. (F) Including a third group, LPS, also revealed a significant (p<0.025) age x group interaction on RLIC FA. Graphs show predicted margins with 95% confidence intervals. Significance values represent corrected p-values for age x group interactions. Abbreviations: RLIC=retrolenticular portion of internal capsule, FA=fractional anisotropy, RD=radial diffusivity, MD= mean diffusivity, AD=axial diffusivity, TD=typically developing, PS= psychosis spectrum, LPS=limited psychosis spectrum.

Secondary analyses of RLIC FA indicated there were significant age x group interactions in both the left (β=-0.0021, F(1,654)=7.72, p=0.0056) and right (β=-0.0024, F(1,647)=9.40, p_ROI_=0.0023) hemispheres. Of the three non-FA diffusivity measures, both RD (β=2.92×10^-6^, F(1,657)=14.21, p=2.00×10^-4^; Figure 3C) and MD (β=7.58×10^-7^, F(1,656)=7.90, p=0.0051; Figure 3D) showed a significant interaction between group and age. TD youth showed a significant decreases in RLIC RD (β=-2.5×10^-6^, p=9.1×10^-13^) and MD (β=-2.9×10^-6^, p=3.4×10^-16^) with increasing age. In PS youth, but there were no significant relationships between age and RLIC RD (β=4.97×10^-7^, p=0.50) or MD (β=-6.8×10^-7^, p=0.39). Neither group showed a significant age x group interaction on AD (Figure 3E).

Inclusion of LPS youth as a third group in the regression model also resulted in a significant age x group interaction (F(2, 692)=5.36, p=0.0049; Figure 3F). Post-hoc contrasts revealed that the slope of the relationship between age and FA for LPS youth did not significantly differ from TD (t(1,692)=-0.02, p=0.982) or PS (t(1,695)=-0.166, p=0.097). Graphical representation of the interaction and evaluation of the regression coefficient between age and RLIC FA in LPS revealed a positive, non-significant association between age and FA (β=0.001).

### Neurocognitive Efficiency Results

PS youth had significantly (p<0.0125) lower efficiency scores in all four cognitive domains: Executive Control, Complex Cognition, Episodic Memory and Social Cognition (Figure 4; Table S3).

**Figure 4.**
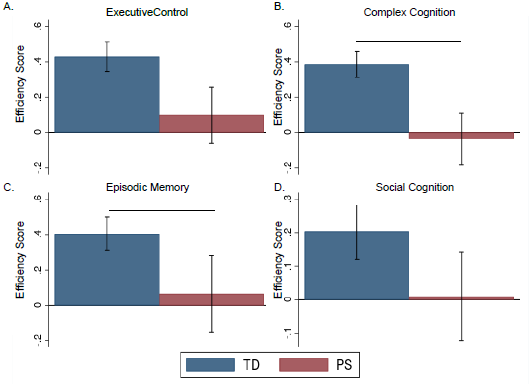
Group differences in cognitive domains. PS youth consistently showed significantly (p<0.0125) lower efficiency scores in all four cognitive domains: (A) Executive Control, (B) Complex Cognition, (C) Episodic Memory and (D) Social Cognition. Efficiency scores reflect the average Z-score (sum of Z-score for accuracy and -1 multiplied by Z-score for speed) per group, such that higher scores indicate better performance. Error bars represent +/- standard error of the group average. Abbreviations: TD=typically developing, PS= psychosis spectrum.

### Post hoc Mediation Analyses

*SLF:* The PS group did not satisfy initial requirements for mediation of the effect of age on any cognitive domain, as both *a* (coefficient for effect of age on SLF) and *b* (coefficient for effect of SLF on cognition) were non-significant (p>0.05) for all domains. However, in the TD group, path coefficients *a* and *b* were significant for Complex Cognition and Episodic Memory, such that scores increased with both age and SLF FA (Table 2), and so mediation was tested. Evaluating the mediation effect in TD with a bootstrapping approach indicated that only the relationship between age and Complex Cognition was significantly mediated by SLF FA, with SLF FA accounting for 27.6% of the total effect between age and Complex Cognition. Path coefficients of mediation analyses for Executive Control and Social Cognition can be found in Table S4.

**Table 2.** Mediation analysis results for TD youth. Abbreviations: TD=typically developing, SLF=superior longitudinal fasciculus, CI=confidence interval.

| Dependent Variable | Path Coefficients (SE) |  |  |  |  | Percent of Total Effect Mediated | Bootstrap 99% CI |
| --- | --- | --- | --- | --- | --- | --- | --- |
|  | age → SLF<br>*a* | SLF → Cognition<br>*b* | Direct Effect<br>*c'* | Total Effect<br>*c=ab+c'* | Mediated Effect<br>*ab* |  |  |
| Complex Cognition<sup>a</sup> | 0.0030**<br>(0.0003) | 6.377**<br>(1.415) | 0.0492**<br>(0.0113) | 0.0683**<br>(0.0107) | 0.0189<sup>c</sup><br>(0.0047) | 27.6 | 0.0066,<br>0.0311 |
| Episodic Memory<sup>b</sup> | 0.0031**<br>(0.0003) | 3.763*<br>(1.85) | 0.0860**<br>(0.0149) | 0.0975**<br>(0.014) | 0.0115<br>(0.0058) | 11.8 | -0.0055,<br>0.0285 |

\* $p<0.05$ , \*\* $p<0.001$ , <sup>c</sup> mediation effect significant at 99% CI
<sup>a</sup> $n=491$ , <sup>b</sup> $n=495$

*RLIC:* In contrast to the SLF, the effect of age on RLIC FA (*a*) was significant for both TD (β=-3.75×10^-6^, F(1,491)=46.18, p=3.13×10^-11^) and PS (β=-3.03×10^-6^, F(1,164)=4.99, p=0.027). However, RLIC FA did not significantly explain any cognitive score in either TD or PS youth (p>0.05). Thus, criteria for the mediation of the relationship between age and each cognitive domain were not met and no further analyses conducted.

## Discussion

Our results provide compelling evidence that subclinical psychotic symptoms are associated with disrupted developmental trajectories of WM in the SLF and RLIC. The SLF is an associative tract with ipsilateral frontoparietal connections [46] that shows pronounced development across adolescence [21, 50, 51] and has a well-established role in fundamental cognitive processes such as working memory, language, and attention [47-49]. The RLIC is a projection tract containing motor and sensory fibers that carry projections from the thalamic pulvinar and lateral geniculate nuclei to association and visual cortices [52]. We have also demonstrated that WM development in the SLF partially mediates the relationship between age and Complex Cognition in TD, but not PS, youth. These findings suggest that alterations in WM neurodevelopment may potentially contribute to cognitive deficits in psychosis.

Of the 25 tracts evaluated, only the SLF and RLIC reached the corrected-threshold for a significant age x group interaction. Consistent with existing studies of WM maturation during typical development [21], we found a linear increase in FA in both the SLF and RLIC in the TD group. The PS group, however, did not show any significant age-related changes in FA in either the SLF or RLIC, suggesting disrupted or blunted maturation of these tracts. Together with findings of age x group interactions for radial but not axial diffusivity, these results collectively suggest that the altered age relationships in PS youth in these tracts may be driven by decreased myelination, rather than axonal degeneration or differences in tract organization [53-55].

Reduced SLF integrity has been reported in both patients with [56] and at risk [57] for SZ, as well as adolescents endorsing subclinical psychotic symptoms [58, 59]. Furthermore, a longitudinal study indicated that high-risk individuals who converted to psychotic disorders had reduced SLF FA at baseline relative to high-risk individuals who did not convert [60]. Our results extend upon these previous findings, suggesting that age-associated alterations of WM development in the SLF as early as late-childhood may be associated with subclinical psychosis and serve as a putative early biomarker for SZ.

Association tracts such as the SLF have previously been related to cognitive function [61-65], and the development of higher-level cognitive processes during adolescence [66, 67] has been linked to increases in FA [30-32]. However, the relationship between the SLF and cognition has been less frequently investigated in the context of youth with psychosis. Furthermore, previous studies have typically evaluated the relationship between cognition and WM separately from that of cognition and age [68], and thus cannot rule out that gains in neurocognitive performance may only relate to FA because of a shared association with age.

Our analysis took the novel approach of testing the degree to which WM integrity mediates development of cognition from childhood through young adulthood, and how this relationship differs in PS. We found a significant positive relationship of age with Complex Cognition and SLF FA in TD, but no such associations in PS youth. Additionally, the association between age and Complex Cognition in TD was significantly mediated by SLF FA, highlighting the tract’s importance in healthy cognitive development. Here, Complex Cognition is a reflection of nonverbal reasoning, language reasoning, and spatial attention [35]. Our finding is consistent with numerous existing studies in which frontoparietal structural connectivity, including the SLF, has been related to attention [69, 70], language [70, 71], spatial working memory [72, 73], and reasoning ability [74] in TD youth. Absence of any significant age-associated changes in Complex Cognition and SLF FA, and thus any corresponding mediation, in PS youth suggests that the presence of subclinical psychotic symptoms disrupts maturation of the SLF which, in turn, affects development of higher-level cognitive function, as demonstrated by the overall significantly lower Complex Cognition observed in PS relative to TD youth. Critically, mediation studies evaluating changes across adulthood into elderly populations found that age-related reductions in SLF FA did *not* mediate corresponding cognitive decline [75, 76], suggesting that its role as a mediator in TD youth may be specific to emergence rather than decline of higher-level cognitive function.

Our findings also revealed an age x group interaction on FA in the RLIC. Reduced FA in the internal capsule of patients with chronic SZ has been consistently reported, with mixed findings in first episode and medication-naïve patients (for review see [23, 77]). However, to our knowledge, there are no studies that focus specifically on the RLIC in relation to psychosis. The strongest previous evidence for a potential association of reduced RLIC FA with psychosis comes from a study in which non-human primates receiving daily exposure to ketamine (an NMDA antagonist used to mimic psychotic symptoms) during adolescence showed reduced FA in multiple tracts, including the RLIC and SLF [78]. Furthermore, a longitudinal study of adults (age 20-40) found that patients with SZ showed an attenuated increase in RLIC FA across a 3-year follow-up relative to controls [79]. Finally, a recent meta-analysis using the same 25 ROIs showed adults with SZ had lower RLIC FA than healthy controls, though the effect size (d=1.2) was not significant [80]. Considered with our findings, these studies may suggest that abnormalities in RLIC FA are most pronounced during the developmental period, with a compensatory increase occurring later in adulthood, or that some other confound in adult patients, such as long-term medication, affects this measure. Further study of this tract in developmental and adult populations is needed to confirm these results.

Projection tracts such as the IC (and here RLIC) are important for corticospinal and sensory communication and have been less frequently investigated in relation to symptomatology of psychiatric disease than associative tracts. Previous studies have postulated that reduced WM in the IC may lead to functional dysconnectivity between the cortex and subcortical structures [81], with corresponding clinically-relevant correlates such as increased positive symptoms [82] and impairments in emotional stability, motivation and executive function [77, 82-84]. Although dysfunctional processing of information has been increasingly implicated in SZ as a putative contributor to impairments in higher-level cognitive function [85], we did not find any significant relationship between RLIC FA and the cognitive measures used here. However, there is evidence that motor- and sensory-related symptoms of psychosis emerge before deficits in higher-order cognition [86]. The RLIC contains both motor and sensory fibers [87], and thus it is possible that RLIC FA mediates age-related changes in these systems rather than cognition.

Finally, our preliminary findings in the LPS group support the hypothesis that the extent of deviation from typical age-associated changes in RLIC and SLF FA is related to the severity of subclinical symptoms. LPS showed an intermediate pattern, with FA values falling between those of TD and PS. This is further evidence that abnormalities in age-associated structural changes are emerging even at the lowest end of the psychosis spectrum.

Strengths of this study include the non-help-seeking, population-based cohort that allows us to eliminate typical confounds of illness such as medication and illness chronicity. Thus, our results are more representative of natural variability than is a typical case-control study. Our results also provide evidence for regional specificity of disruptions in FA development, and expand upon these findings by also evaluating non-FA measures. Finally, there is a paucity of studies examining how regional WM measures may mediate the relationship between age and other behavioral and cognitive endophenotypes of SZ. This is likely attributable to the rarity of access to both neuroimaging and cognitive data for a sample size of this magnitude that spans such a broad age range. Future studies incorporating more precise measures of development and maturation, such as pubertal stage, could help to further refine our understanding of the developmental relationships reported here. Additionally, prospective longitudinal data will be essential to determine neural biomarkers that predict subsequent outcome.

## Conclusions

In this population-based young sample, youth with PS symptoms showed disrupted development of fronto–parietal WM connectivity, and — unlike in TD youth — did not show mediation of the relationship between age and cognition by FA in this tract. Collectively, these findings suggest disturbances in the developmental trajectories of the SLF and RLIC may be an early biomarker for psychosis and that the SLF may contribute to cognitive deficits. Future investigations should pursue longitudinal investigations to determine their predictive validity, as well as pre-clinical experimental studies to provide mechanistic explanations and potentially identify critical temporal and anatomical targets for neurotherapeutic interventions.

## Acknowledgments

This research was supported by National Institute of Mental Health (NIMH) grant R01 MH107250 awarded to CEB and NIMH grant R01 MH101506 awarded to KHK. We would like to acknowledge that all data were obtained via the Database of Genotypes and Phenotypes (dbGap) platform (Project #6984, Karlsgodt, Releases 1 and 2).

## References

1. Moreno-Küstner, B., C. Martín, and L. Pastor, Prevalence of psychotic disorders and its association with methodological issues. A systematic review and meta-analyses. PLoS One, 2018. 13(4): p. e0195687.

2. Strauss, J.S., Hallucinations and delusions as points on continua function. Rating scale evidence. Arch Gen Psychiatry, 1969. 21(5): p. 581-6.

3. DeRosse, P. and K.H. Karlsgodt, Examining the Psychosis Continuum. Curr Behav Neurosci Rep, 2015. 2(2): p. 80-89.

4. Linscott, R.J. and J. van Os, An updated and conservative systematic review and meta-analysis of epidemiological evidence on psychotic experiences in children and adults: on the pathway from proneness to persistence to dimensional expression across mental disorders. Psychol Med, 2013. 43(6): p. 1133-49.

5. van Os, J., et al., A systematic review and meta-analysis of the psychosis continuum: evidence for a psychosis proneness-persistence-impairment model of psychotic disorder. Psychol Med, 2009. 39(2): p. 179-95.

6. Koyanagi, A., Psychotic-like experiences and happiness in the English general population. J Affect Disord, 2017. 222: p. 211-217.

7. Jang, J.H., et al., Psychotic-like experiences and their relationship to suicidal ideation in adolescents. Psychiatry Res, 2014. 215(3): p. 641-5.

8. Fisher, H.L., et al., Specificity of childhood psychotic symptoms for predicting schizophrenia by 38 years of age: a birth cohort study. Psychol Med, 2013. 43(10): p. 2077-86.

9. Correll, C.U., et al., Research in people with psychosis risk syndrome: a review of the current evidence and future directions. J Child Psychol Psychiatry, 2010. 51(4): p. 390-431.

10. Fusar-Poli, P., et al., Predicting psychosis: meta-analysis of transition outcomes in individuals at high clinical risk. Arch Gen Psychiatry, 2012. 69(3): p. 220-9.

11. David, A.S. and O. Ajnakina, Psychosis as a continuous phenotype in the general population: the thin line between normality and pathology. World Psychiatry, 2016. 15(2): p. 129-30.

12. Poulton, R., et al., Children’s self-reported psychotic symptoms and adult schizophreniform disorder: a 15-year longitudinal study. Arch Gen Psychiatry, 2000. 57(11): p. 1053-8.

13. Dominguez, M.D., et al., Evidence that onset of clinical psychosis is an outcome of progressively more persistent subclinical psychotic experiences: an 8-year cohort study. Schizophr Bull, 2011. 37(1): p. 84-93.

14. Escher, S., et al., Independent course of childhood auditory hallucinations: a sequential 3-year follow-up study. Br J Psychiatry Suppl, 2002. 43: p. s10-8.

15. Rössler, W., et al., Psychotic experiences in the general population: a twenty-year prospective community study. Schizophr Res, 2007. 92(1-3): p. 1-14.

16. Lenroot, R.K., et al., Sexual dimorphism of brain developmental trajectories during childhood and adolescence. Neuroimage, 2007. 36(4): p. 1065-73.

17. Giedd, J.N., et al., Anatomical brain magnetic resonance imaging of typically developing children and adolescents. J Am Acad Child Adolesc Psychiatry, 2009. 48(5): p. 465-70.

18. Tamnes, C.K., et al., Brain maturation in adolescence and young adulthood: regional age-related changes in cortical thickness and white matter volume and microstructure. Cereb Cortex, 2010. 20(3): p. 534-48.

19. Thomason, M.E. and P.M. Thompson, Diffusion imaging, white matter, and psychopathology. Annu Rev Clin Psychol, 2011. 7: p. 63-85.

20. Lebel, C., et al., Microstructural maturation of the human brain from childhood to adulthood. Neuroimage, 2008. 40(3): p. 1044-55.

21. Peters, B.D., et al., White matter development in adolescence: diffusion tensor imaging and meta-analytic results. Schizophr Bull, 2012. 38(6): p. 1308-17.

22. Pettersson-Yeo, W., et al., Dysconnectivity in schizophrenia: where are we now? Neurosci Biobehav Rev, 2011. 35(5): p. 1110-24.

23. Wheeler, A.L. and A.N. Voineskos, A review of structural neuroimaging in schizophrenia: from connectivity to connectomics. Front Hum Neurosci, 2014. 8: p. 653.

24. Satterthwaite, T.D., et al., Structural Brain Abnormalities in Youth With Psychosis Spectrum Symptoms. JAMA Psychiatry, 2016. 73(5): p. 515-24.

25. Krabbendam, L., et al., Familial covariation of the subclinical psychosis phenotype and verbal fluency in the general population. Schizophr Res, 2005. 74(1): p. 37-41.

26. Ziermans, T.B., Working memory capacity and psychotic-like experiences in a general population sample of adolescents and young adults. Front Psychiatry, 2013. 4: p. 161.

27. Kelleher, I., et al., Neurocognition in the extended psychosis phenotype: performance of a community sample of adolescents with psychotic symptoms on the MATRICS neurocognitive battery. Schizophr Bull, 2013. 39(5): p. 1018-26.

28. Martín-Santiago, O., et al., [Relationship between subclinical psychotic symptoms and cognitive performance in the general population]. Rev Psiquiatr Salud Ment, 2016. 9(2): p. 78-86.

29. Barnett, J.H., V. Hachinski, and A.D. Blackwell, Cognitive health begins at conception: addressing dementia as a lifelong and preventable condition. BMC Med, 2013. 11: p. 246.

30. Moseley, M., R. Bammer, and J. Illes, Diffusion-tensor imaging of cognitive performance. Brain Cogn, 2002. 50(3): p. 396-413.

31. Schmithorst, V.J., et al., Cognitive functions correlate with white matter architecture in a normal pediatric population: a diffusion tensor MRI study. Hum Brain Mapp, 2005. 26(2): p. 139-47.

32. Mabbott, D.J., et al., White matter growth as a mechanism of cognitive development in children. Neuroimage, 2006. 33(3): p. 936-46.

33. Satterthwaite, T.D., et al., The Philadelphia Neurodevelopmental Cohort: A publicly available resource for the study of normal and abnormal brain development in youth. Neuroimage, 2016. 124(Pt B): p. 1115-9.

34. Calkins, M.E., et al., The psychosis spectrum in a young U.S. community sample: findings from the Philadelphia Neurodevelopmental Cohort. World Psychiatry, 2014. 13(3): p. 296-305.

35. Moore, T.M., et al., Psychometric properties of the Penn Computerized Neurocognitive Battery. Neuropsychology, 2015. 29(2): p. 235-46.

36. Smith, S.M., et al., Tract-based spatial statistics: voxelwise analysis of multi-subject diffusion data. Neuroimage, 2006. 31(4): p. 1487-505.

37. Jahanshad, N., et al., Multi-site genetic analysis of diffusion images and voxelwise heritability analysis: a pilot project of the ENIGMA-DTI working group. Neuroimage, 2013. 81: p. 455-69.

38. Mori, S., et al., MRI Atlas of Human White Matter. 2005, Elsevier Science. p. 276.

39. Wakana, S., et al., Reproducibility of quantitative tractography methods applied to cerebral white matter. Neuroimage, 2007. 36(3): p. 630-44.

40. Hua, K., et al., Tract probability maps in stereotaxic spaces: analyses of white matter anatomy and tract-specific quantification. Neuroimage, 2008. 39(1): p. 336-47.

41. Alexander, A.L., et al., Diffusion tensor imaging of the brain. Neurotherapeutics, 2007. 4(3): p. 316-29.

42. Basser, P.J. and D.K. Jones, Diffusion-tensor MRI: theory, experimental design and data analysis - a technical review. NMR Biomed, 2002. 15(7-8): p. 456-67.

43. Chang, E.H., et al., Diffusion tensor imaging measures of white matter compared to myelin basic protein immunofluorescence in tissue cleared intact brains. Data Brief, 2017. 10: p. 438-443.

44. MacKinnon, D.P., A.J. Fairchild, and M.S. Fritz, Mediation analysis. Annu Rev Psychol, 2007. 58: p. 593-614.

45. DP, M. and D. JH, Estimation of mediated effects in prevention studies. Eval Rev., 1993. 17(144-58).

46. Schmahmann, J.D., et al., Cerebral white matter: neuroanatomy, clinical neurology, and neurobehavioral correlates. Ann N Y Acad Sci, 2008. 1142: p. 266-309.

47. Mesulam, M.M., From sensation to cognition. Brain, 1998. 121 (Pt 6): p. 1013-52.

48. Petrides, M. and D.N. Pandya, Comparative cytoarchitectonic analysis of the human and the macaque ventrolateral prefrontal cortex and corticocortical connection patterns in the monkey. Eur J Neurosci, 2002. 16(2): p. 291-310.

49. Madhavan, K.M., et al., Superior longitudinal fasciculus and language functioning in healthy aging. Brain Res, 2014. 1562: p. 11-22.

50. Lebel, C. and C. Beaulieu, Longitudinal development of human brain wiring continues from childhood into adulthood. J Neurosci, 2011. 31(30): p. 10937-47.

51. Giorgio, A., et al., Longitudinal changes in grey and white matter during adolescence. Neuroimage, 2010. 49(1): p. 94-103.

52. Larry, R., et al., Fundamental Neuroscience. 4th Edition ed. 2012, Waltham, MA: Elsevier.

53. Beaulieu, C. and P.S. Allen, Determinants of anisotropic water diffusion in nerves. Magn Reson Med, 1994. 31(4): p. 394-400.

54. Song, S.K., et al., Diffusion tensor imaging detects and differentiates axon and myelin degeneration in mouse optic nerve after retinal ischemia. Neuroimage, 2003. 20(3): p. 1714-22.

55. Wozniak, J.R. and K.O. Lim, Advances in white matter imaging: a review of in vivo magnetic resonance methodologies and their applicability to the study of development and aging. Neurosci Biobehav Rev, 2006. 30(6): p. 762-74.

56. Karlsgodt, K.H., et al., Diffusion tensor imaging of the superior longitudinal fasciculus and working memory in recent-onset schizophrenia. Biol Psychiatry, 2008. 63(5): p. 512-8.

57. Karlsgodt, K.H., et al., White matter integrity and prediction of social and role functioning in subjects at ultra-high risk for psychosis. Biol Psychiatry, 2009. 66(6): p. 562-9.

58. O’Hanlon, E., et al., White Matter Differences Among Adolescents Reporting Psychotic Experiences: A Population-Based Diffusion Magnetic Resonance Imaging Study. JAMA Psychiatry, 2015. 72(7): p. 668-77.

59. DeRosse, P., et al., White Matter Abnormalities Associated With Subsyndromal Psychotic-Like Symptoms Predict Later Social Competence in Children and Adolescents. Schizophr Bull, 2017. 43(1): p. 152-159.

60. Bloemen, O.J., et al., White-matter markers for psychosis in a prospective ultra-high-risk cohort. Psychol Med, 2010. 40(8): p. 1297-304.

61. Felten, D.L., M.K. O’Banion, and M.S. Maida, Telencephalon, in Netter’s Atlas of Neuroscience. 2016, Elsevier Inc. p. 463-477.

62. Bai, F., et al., Abnormal integrity of association fiber tracts in amnestic mild cognitive impairment. J Neurol Sci, 2009. 278(1-2): p. 102-6.

63. Gold, B.T., et al., Speed of lexical decision correlates with diffusion anisotropy in left parietal and frontal white matter: evidence from diffusion tensor imaging. Neuropsychologia, 2007. 45(11): p. 2439-46.

64. Potapov, A.A., et al., [The long-associative pathway of the white matter: modern view from the perspective of neuroscience]. Zh Vopr Neirokhir Im N N Burdenko, 2014. 78(5): p. 66-77; discussion 77.

65. Turken, A., et al., Cognitive processing speed and the structure of white matter pathways: convergent evidence from normal variation and lesion studies. Neuroimage, 2008. 42(2): p. 1032-44.

66. Choudhury, S., S.J. Blakemore, and T. Charman, Social cognitive development during adolescence. Soc Cogn Affect Neurosci, 2006. 1(3): p. 165-74.

67. Luna, B., Developmental changes in cognitive control through adolescence. Adv Child Dev Behav, 2009. 37: p. 233-78.

68. Madden, D.J., I.J. Bennett, and A.W. Song, Cerebral white matter integrity and cognitive aging: contributions from diffusion tensor imaging. Neuropsychol Rev, 2009. 19(4): p. 415-35.

69. Hamilton, L.S., et al., Reduced white matter integrity in attention-deficit hyperactivity disorder. Neuroreport, 2008. 19(17): p. 1705-8.

70. Urger, S.E., et al., The superior longitudinal fasciculus in typically developing children and adolescents: diffusion tensor imaging and neuropsychological correlates. J Child Neurol, 2015. 30(1): p. 9-20.

71. Brauer, J., A. Anwander, and A.D. Friederici, Neuroanatomical prerequisites for language functions in the maturing brain. Cereb Cortex, 2011. 21(2): p. 459-66.

72. Nagy, Z., H. Westerberg, and T. Klingberg, Maturation of white matter is associated with the development of cognitive functions during childhood. J Cogn Neurosci, 2004. 16(7): p. 1227-33.

73. Vestergaard, M., et al., White matter microstructure in superior longitudinal fasciculus associated with spatial working memory performance in children. J Cogn Neurosci, 2011. 23(9): p. 2135-46.

74. Wendelken, C., et al., Frontoparietal Structural Connectivity in Childhood Predicts Development of Functional Connectivity and Reasoning Ability: A Large-Scale Longitudinal Investigation. J Neurosci, 2017. 37(35): p. 8549-8558.

75. Madden, D.J., et al., Cerebral white matter integrity mediates adult age differences in cognitive performance. J Cogn Neurosci, 2009. 21(2): p. 289-302.

76. Salami, A., et al., Age-related white matter microstructural differences partly mediate age-related decline in processing speed but not cognition. Biochim Biophys Acta, 2012. 1822(3): p. 408-15.

77. Vitolo, E., et al., White matter and schizophrenia: A meta-analysis of voxel-based morphometry and diffusion tensor imaging studies. Psychiatry Res Neuroimaging, 2017. 270: p. 8-21.

78. Li, Q., et al., Chronic Ketamine Exposure Causes White Matter Microstructural Abnormalities in Adolescent Cynomolgus Monkeys. Front Neurosci, 2017. 11: p. 285.

79. Domen, P., et al., Differential Time Course of Microstructural White Matter in Patients With Psychotic Disorder and Individuals at Risk: A 3-Year Follow-up Study. Schizophr Bull, 2017. 43(1): p. 160-170.

80. Kelly, S., et al., Widespread white matter microstructural differences in schizophrenia across 4322 individuals: results from the ENIGMA Schizophrenia DTI Working Group. Mol Psychiatry, 2018. 23(5): p. 1261-1269.

81. Kubicki, M., et al., DTI and MTR abnormalities in schizophrenia: analysis of white matter integrity. Neuroimage, 2005. 26(4): p. 1109-18.

82. Yao, L., et al., Association of white matter deficits with clinical symptoms in antipsychotic-naive first-episode schizophrenia: an optimized VBM study using 3T. MAGMA, 2014. 27(4): p. 283-90.

83. Guo, W., et al., Right lateralized white matter abnormalities in first-episode, drug-naive paranoid schizophrenia. Neurosci Lett, 2012. 531(1): p. 5-9.

84. Lyu, H., et al., Regional white matter abnormalities in drug-naive, first-episode schizophrenia patients and their healthy unaffected siblings. Aust N Z J Psychiatry, 2015. 49(3): p. 246-54.

85. Leitman, D.I., et al., Sensory deficits and distributed hierarchical dysfunction in schizophrenia. Am J Psychiatry, 2010. 167(7): p. 818-27.

86. Forsyth, J.K. and D.A. Lewis, Mapping the Consequences of Impaired Synaptic Plasticity in Schizophrenia through Development: An Integrative Model for Diverse Clinical Features. Trends Cogn Sci, 2017. 21(10): p. 760-778.

87. Ribas, G.C., Applied Cranial-Cerebral Anatomy: Brain Architecture and Anatomically Oriented Microneurosurgery. 2018, Cambridge University Press: Cambridge, United Kingdom. p. 15-61.

